# Omics BioAnalytics: Reproducible Research using R Shiny and Alexa

**DOI:** 10.1101/2020.05.05.024323

**Authors:** Amrit Singh, Scott J. Tebbutt, Bruce M. McManus

**Affiliations:** Centre of Excellence for Prevention of Organ Failure, Vancouver, British Columbia, Canada; Department of Pathology and Laboratory Medicine, University of British Columbia, Vancouver, British Columbia, Canada; Department of Medicine, University of British Columbia, Vancouver, British Columbia, Canada

## Abstract

**Summary:** High-throughput technologies produce complex high-dimensional datasets which are analyzed using a variety of ever-evolving bioinformatics tools. Well-designed web frameworks enable more intuitive and efficient analysis such that less time is spent on coding and more time is spent on interpretation of results and addressing insightful biological questions aided by interactive visualizations. Here, we present Omics BioAnalytics, a full-service Web framework that enables comprehensive, multi-level characterization, analysis, and integration of omics datasets. Blending web-based (R Shiny) and voice-based (Amazon’s Alexa) analytics, Omics BioAnalytics can be used both by expert computational biologists and non-coding biological domain experts, alike. Our web framework can be utilized to explore complex datasets and identify biosignatures and discriminative biomarkers of health and disease processes, and generate testable hypotheses relating to underlying molecular mechanisms.

**Availability:** Omics BioAnalytics is freely available at https://github.com/singha53/omicsBioAnalytics and the web app is deployed at https://amritsingh.shinyapps.io/omicsBioAnalytics/. The source code for the companion Alexa Skill can be found at https://github.com/singha53/omics-bioanalytics-alexa-skill.

## Introduction

Data wrangling (preprocessing and normalization) of high-dimensional omic datasets are specific to different high-throughput technologies such as microarrays, sequencing, and mass spectrometry. However, the downstream analytical methods, including those applied for exploratory data analysis, differential expression analysis, pathway analysis for biological enrichment, and biomarker discovery using machine learning algorithms, are often agnostic to the original measurement technology and utilize extensive bioinformatics pipelines. Recent advances in web framework technologies have allowed for the analysis of complex datasets by users with little programming experience. The R Shiny framework is being heavily leveraged in developing interactive web apps for high-throughput omics data (*e.g*. WIIsON (Schultheis *et al*., 2019), GENAVi (Reyes *et al*., 2019), ShinyOmics (Surujon and van Opijnen, 2020)). Here, we present Omics BioAnalytics (Fig. 1), extending existing work in a number of innovative and/or useful ways, including: 1) descriptive statistics and statistical inference of metadata (*e.g*. demographics, clinical data, phenotypes, sample labels); 2) analyses of multiple omics (multi-omics) datasets with associated metadata 3) greater number of analyses that can be performed such as drug- and pathway-enrichment analyses and ensemble classification models of multi-omics data; and 4) dynamic reporting for generating customized documents. Importantly, we have created a complementary multi-modal Alexa Skill which allows for voice-based analytics and visualization of multi-omics data. These practices will enable reproducible workflows for omics-based biomedical and other life-science studies from data input to report generation.

**Fig. 1.**
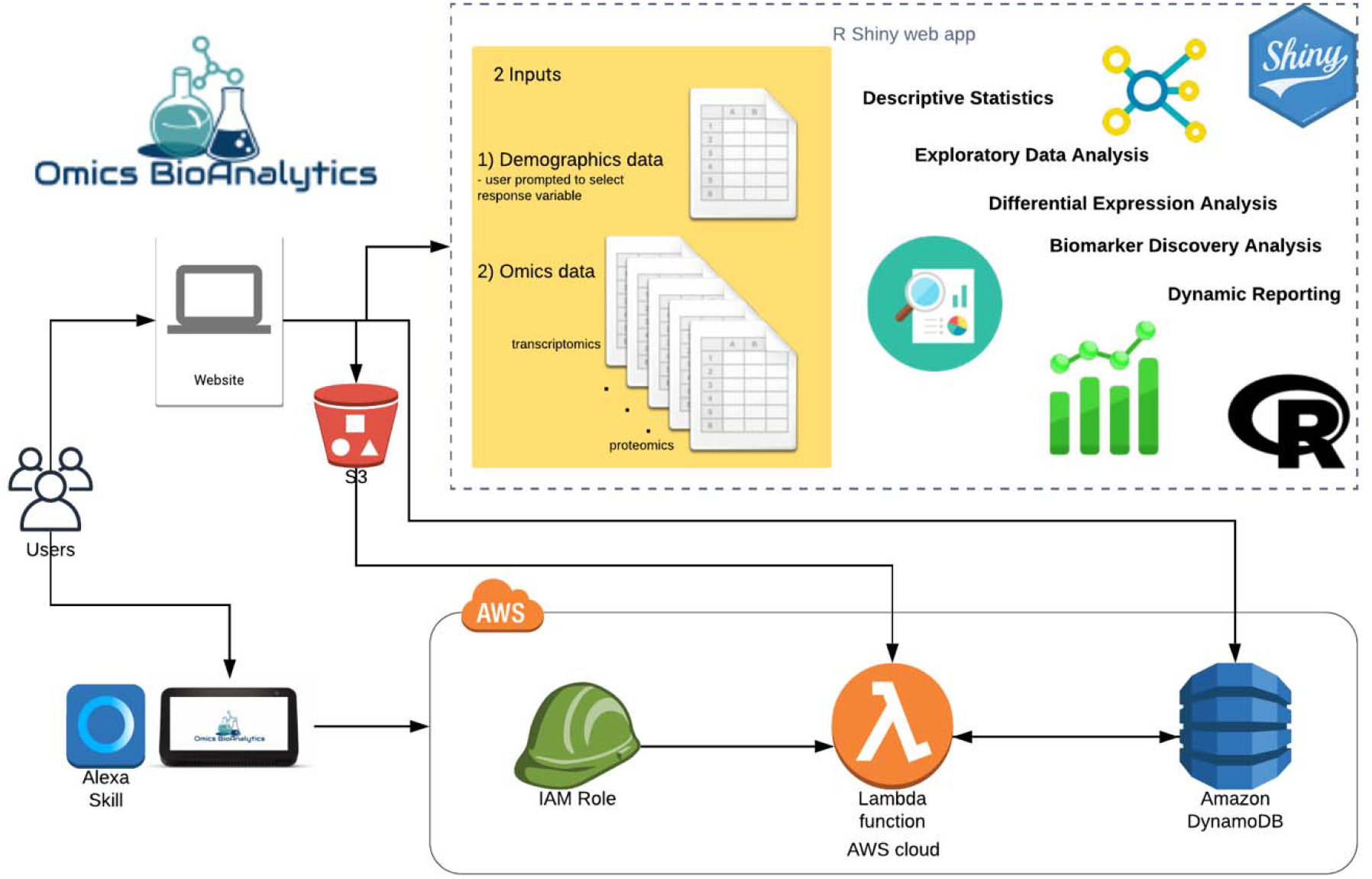
Overview of Omics BioAnalytics. The web app allows users to upload multi-omics data collected on the same set of subjects (samples) as well as their associated demographic variables (phenotypic data). The user can than perform various analyses such as exploratory data analysis, differential expression analysis and biomarker discovery analysis using the web app, or multimodal Alexa Skill. Using the web app, users can also create reports of their analyses.

## Methods

### Data Format

Metadata including demographic variables (*p*) such as age and sex are commonly found along the columns of a spreadsheet whereas the individual observations (*n*) are found along the rows, *i.e. n x p*. The reverse is generally true for omics data, *i.e. p x n*. Furthermore, different bioinformatics tools work with different data orientations, for example *limma* (Ritchie *et al*., 2015) (tool for differential expression) expects a *p x n* format, whereas *glmnet (Zou and Hastie, 2005)* (tool for developing supervised models) expects a *n x p* format. Omics BioAnalytics assumes that the metadata file and arbitrary number of omics data files are in a *n x p* format. The user uploads a metadata file and is than prompted to select a response variable (categorical variable present in the metadata file) and a reference level used as the baseline category. The user can then upload multiple omics data files and click the Run Analysis button.

### Web-based Analytics

#### Descriptive Statistics and Statistical Inference of Metadata

Biological samples (*e.g*. animals, cell lines) have associated metadata (*e.g*. demographics, clinical variables, phenotypic data, sample information) such as age, sex, heart rate, batch information, and disease status. Investigators are often interested in stratifying the data based on some phenotypic/response variable such as disease status. Therefore, we compile basic descriptive statistics as well as conduct hypothesis-testing for variables in the metadata file with respect to the specified response variable. This includes testing for linear regression assumptions (Peña and Slate, 2006) and performing parametric/non-parametric tests for response variables with two or more categories.

#### Exploratory Data Analysis

For each omic dataset provided, a separate dashboard is dynamically generated to display the results of Principal Component Analysis (PCA). The *k* principal components are depicted using a scatter plot matrix and colour-coded by the response variable, where the value of *k* is specified by the user. A heatmap depicts whether there is a pairwise association between each demographic variable and principal component in order to attribute the major sources of variation in each omic data to known sample labels (*e.g*. batch, response).

#### Differential Expression Analysis

For each omic dataset provided, a separate dashboard is dynamically generated to display the results of the differential expression analysis, including an interactive volcano plot [-log_10_(P-value) vs. log_2_(fold-change)] that specifies the significant and not significant variables based on a user controlled false discovery rate (FDR) interactive slider. Clicking on the points of the volcano plot displays each individual variable for closer interrogation. Since the volcano plot only displays pairwise comparisons, for response variables with multiple categories, the user can select which pairwise comparison to view as well as the type of methodology to use including ordinary least squares (OLS), *limma* (suited for microarrays) and *limma voom* (suited for RNA-Seq data) (Ritchie *et al*., 2015). For omics datasets in which gene symbols are detected (*e.g*. transcriptomics, proteomics), The up- and down-regulated significant variables are used to determine biologically enriched pathways (depicted using interactive dot plots) and drug candidates that can be used to reverse the expression profiles (depicted using interactive heatmaps), using the EnrichR API (Kuleshov *et al*., 2016).

#### Biomarker Discovery Analysis

To identify biomarker panels consisting of a subset of variables from each omic dataset, penalized regression models (Zou and Hastie, 2005) are developed for each user-specified omics dataset and the ensemble of multiple omics datasets. Repeated cross-validation is used to determine the optimal values for hyperparameters based on a user-defined grid. The performance of the optimal single and ensemble biomarker panels is depicted using receiver operating characteristic curves. The importance of selected omic biomarkers are displayed using interactive dot plots and overlaps between the single and ensemble biomarker panels are shown using an intersection plot. PCA component plots and heatmaps are used to depict the clustering of the samples based on the selected biomarkers in single and ensemble biomarker panels. The variables of the ensemble biomarker panel are depicted using a correlation network (cut-off controlled by user) and overlaid with the results of pathway enrichment analysis (FDR controlled by user) in order to attribute non-curated variables with variables that are members of curated pathway databases.

#### Dynamic Reporting

For each analysis, figures and tables can be downloaded by the user. Similar to a manuscript, different sections can be created using markdown syntax, and figures and tables can be attached to these sections. This process results in many HTML elements that can be rearranged by drag and drop. Finally, a Word document can be generated consisting of the specified headings, text, figures, and tables for dissemination.

### Voice-based Analytics

Voice-based analytics are useful in querying key performance indices (KPIs) in real-time using technologies such as Amazon Alexa and Google Nest. This can easily be extended to bioinformatics data analysis, as KPIs can include the number of samples, number of features in each omics dataset, list of significant variables and pathways which can be stored in unstructured databases such as Amazon DynamoDB [a NoSQL database from Amazon Web Services (AWS)] and figures can be stored in Amazon Simple Storage Service (S3). The Multi-modal Alexa Skill is then a combination of an interaction model (consisting of user utterances) and an AWS Lambda function (handler function that responds to user utterances). Upon uploading the data files on the Omics BioAnalytics web-app, the user can click the Alexa button to perform voice-based analytics using the Omics BioAnalytics Alexa Skill. The web-app generates a specific id for the analysis, which the user can use to retrieve the analysis when using the Alexa Skill. The analysis is performed by the web-app and the KPIs and figures are stored on DynamoDB and S3, respectively. The lambda function will retrieve the appropriate KPIs and figures to display on the Alexa device based on the user’s voice commands.

### Application

Using public datasets we used our application to: 1) identify compounds that reverse the gene expression patterns observed in lung cells treated with SARS-COV-2 (Blanco-Melo *et al*., 2020) and 2) identify multi-omic biomarker panels of heart failure hospitalizations (Singh *et al*., 2019) (see Supplementary Information).

## Discussion and Conclusion

Omics BioAnalytics is an initiative to reduce the barrier of entry for non-specialists in bioinformatics using both web- and voice-based analytics. The use of dynamic dashboards, dynamic reporting, interactive visualizations and voice-enabled analytics using various bioinformatics tools will enable and expedite a thorough interrogation of omics datasets.

