## Supplementary Information for "Omics BioAnalytics: Reproducible Research using R Shiny and Alexa"

All of the following analyses were performed at <https://amritsingh.shinyapps.io/omicsBioAnalytics/>. Links to download the associated datasets are also provided on the web application.

**Case Study 1: Identify compounds that reverse the gene expression patterns observed in lung cells treated with SARS-COV-2**

*Methods:* RNA-Sequencing was performed on RNA collected from lung epithelial cells under control and infected conditions (20 samples) ([**GSE147507**](https://www.ncbi.nlm.nih.gov/geo/query/acc.cgi?acc=GSE147507)) (Blanco-Melo *et al.*, 2020). Data prefiltering was performed to remove gene transcripts with zero variance and low abundance resulting in 10,563 gene transcripts (<https://github.com/singha53/omicsBioAnalytics/blob/master/inst/extdata/covid19/covid19.md>).

Steps:

1. **Data upload**

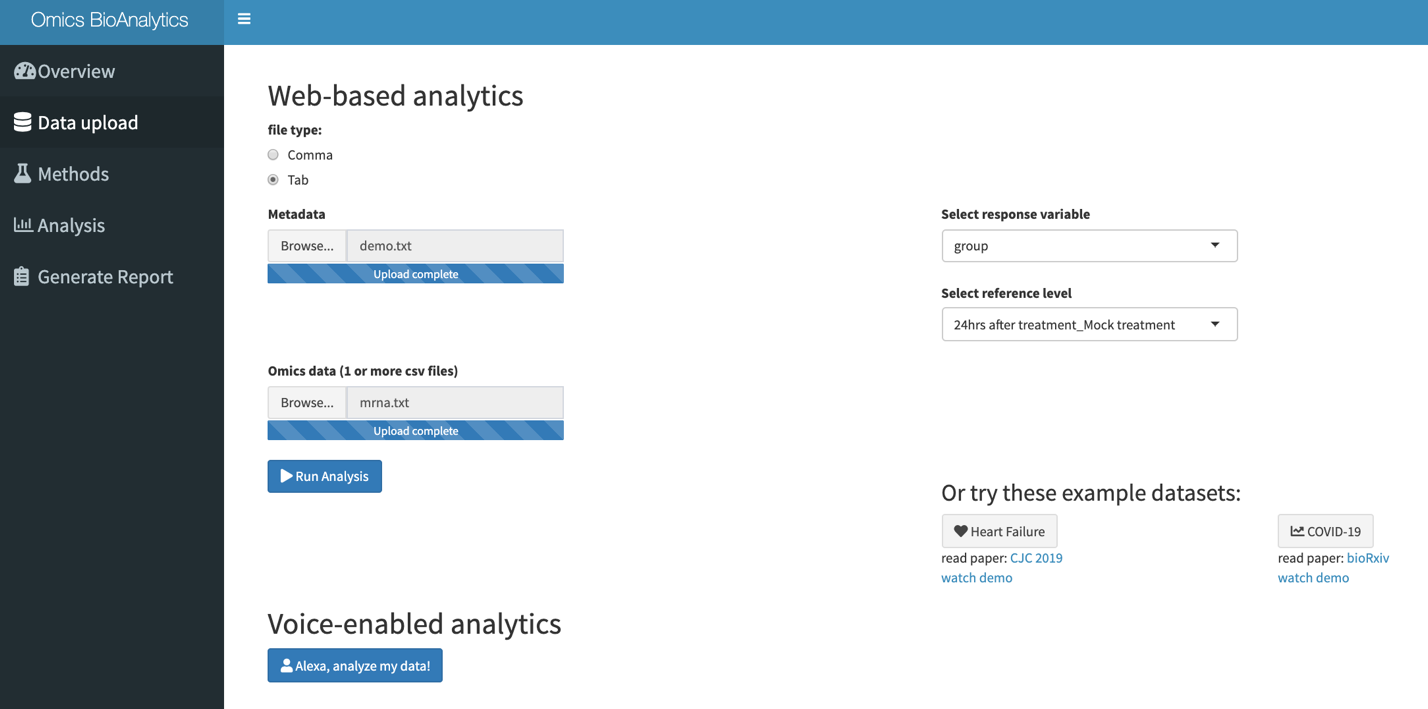

**Supplementary Figure 1. Data upload of COVID-19 datasets.** Metadata of the samples can be uploaded which prompts the user to select a response variable as well as a reference category. In this case, the response variable *group* and the reference category *24hrs after treatment_Mock treatment* were selected. One or more omics datasets can also be uploaded, in this case one text file named *mrna* was uploaded. All files have samples along the rows and variables along the columns.

2. **Differential Expression Analysis**

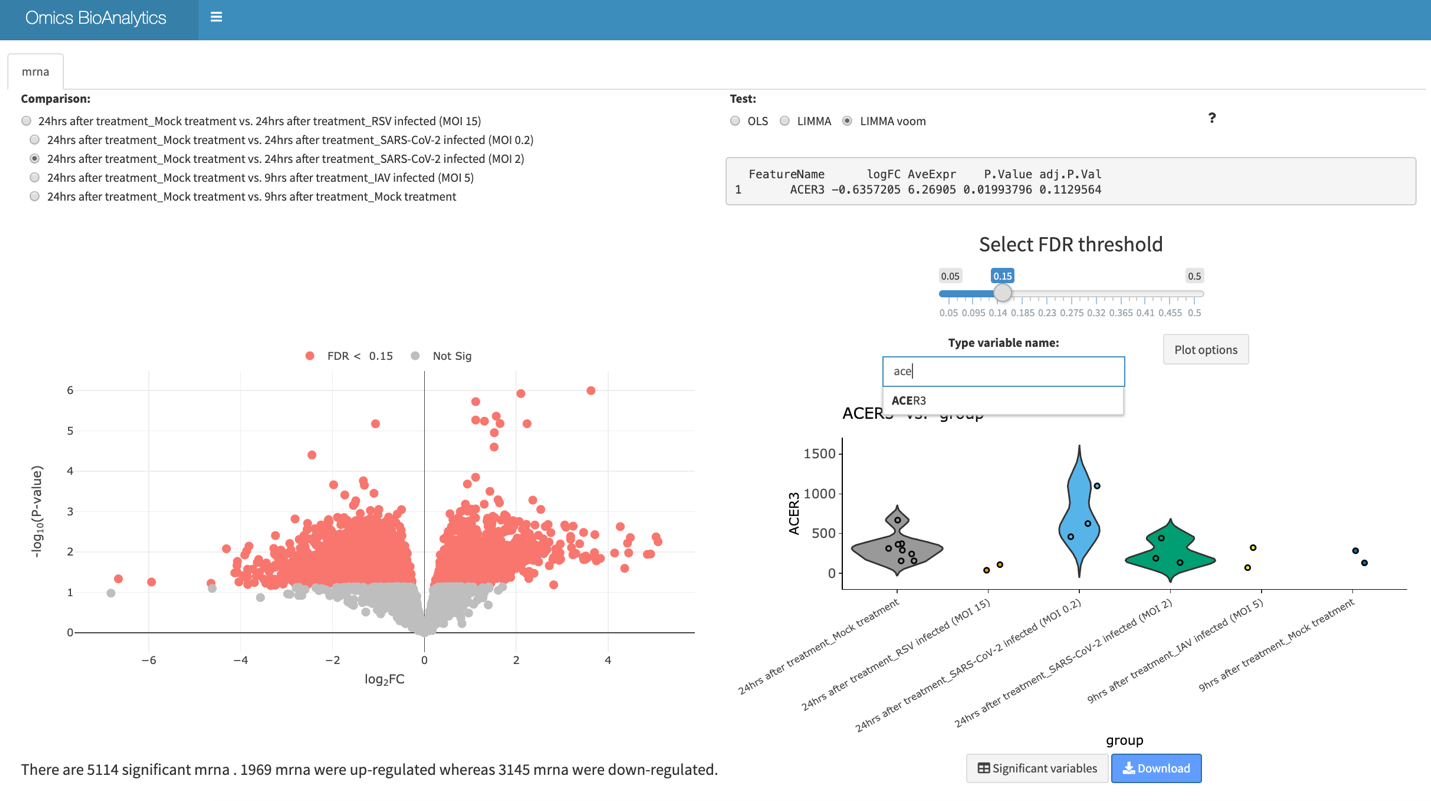

**Supplementary Figure 2: Differential expression analysis of SARS-COV-2 RNA-Seq data.** 5114 differentially expressed genes (mRNA) were identified comparing mock treatment (24h) with SARS-COV-2 infected cells at an FDR=15%. All possible pairwise comparisons can be conducted by using the radio buttons. Clicking on points on the volcano plot or typing in the name of gene (with autocompletion of variables that exist in the current dataset) can be used to plot the expression counts of a single variable, in this case, *ACER3*. LIMMA voom which is suited for RNA-Seq count data was used for this analysis. Users can see the list of significant variables by clicking on the *Significant variable* button or download the results for their own purposes. Plots can be customized and downloaded.

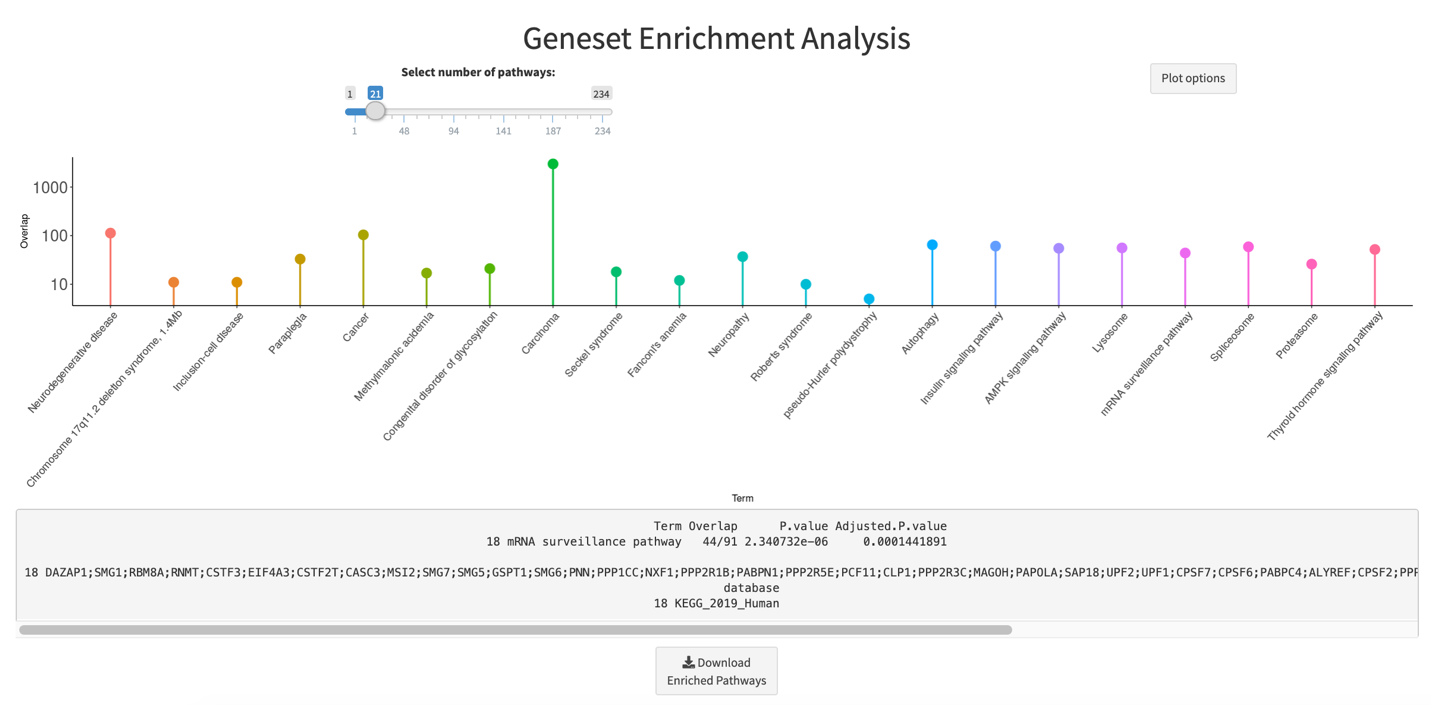

**Supplementary Figure 3: Geneset enrichment analysis using EnrichR.** For datasets in which the web application detects gene symbols, gene set enrichment is automatically performed using the EnrichR API (Kuleshov *et al.*, 2016) using the Jensen_DISEASES, KEGG_2019_Human and WikiPathways_2019 gene set collections based on the list of significant variables from Figure 2. 21 of the 234 significant pathways (FDR=15%) are plotted for easy of viewing (see slider). Clicking on the tip of each point provides additional information such as the overlap between members of the significant gene list and total number of genes in the gene set, p-value, adjusted p-value, and overlapping significant gene symbols. This data can be download in a table form by clicking on the *Download Enriched Pathways* button.

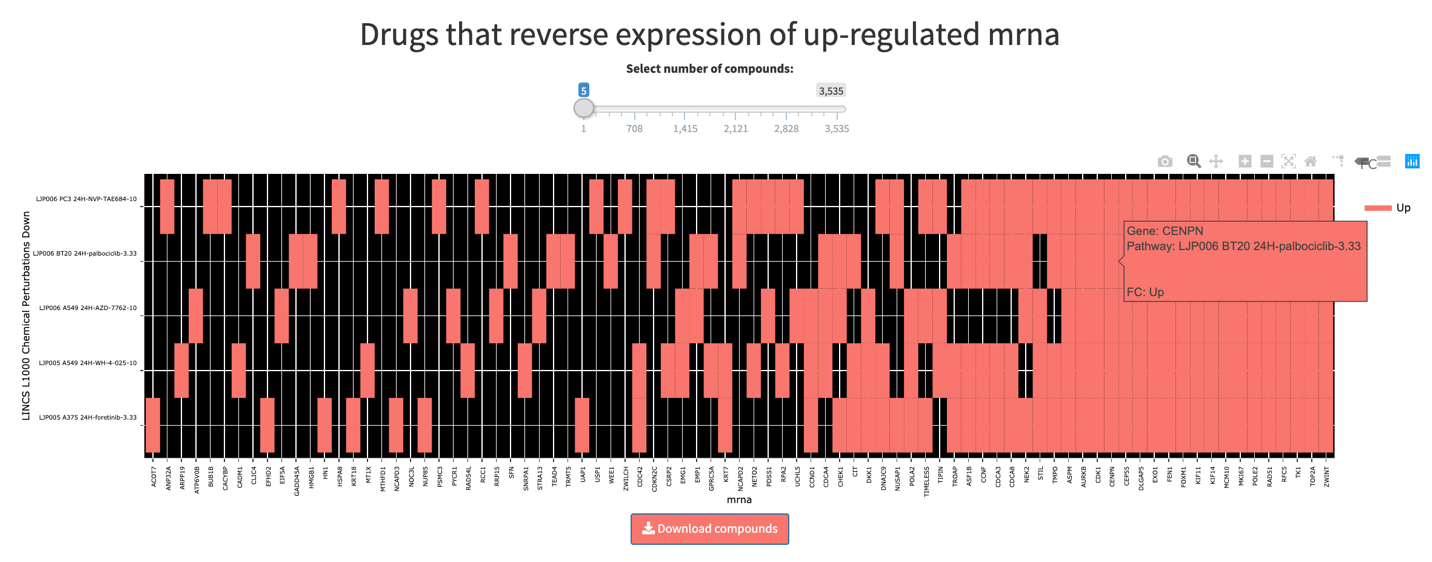

**A.**

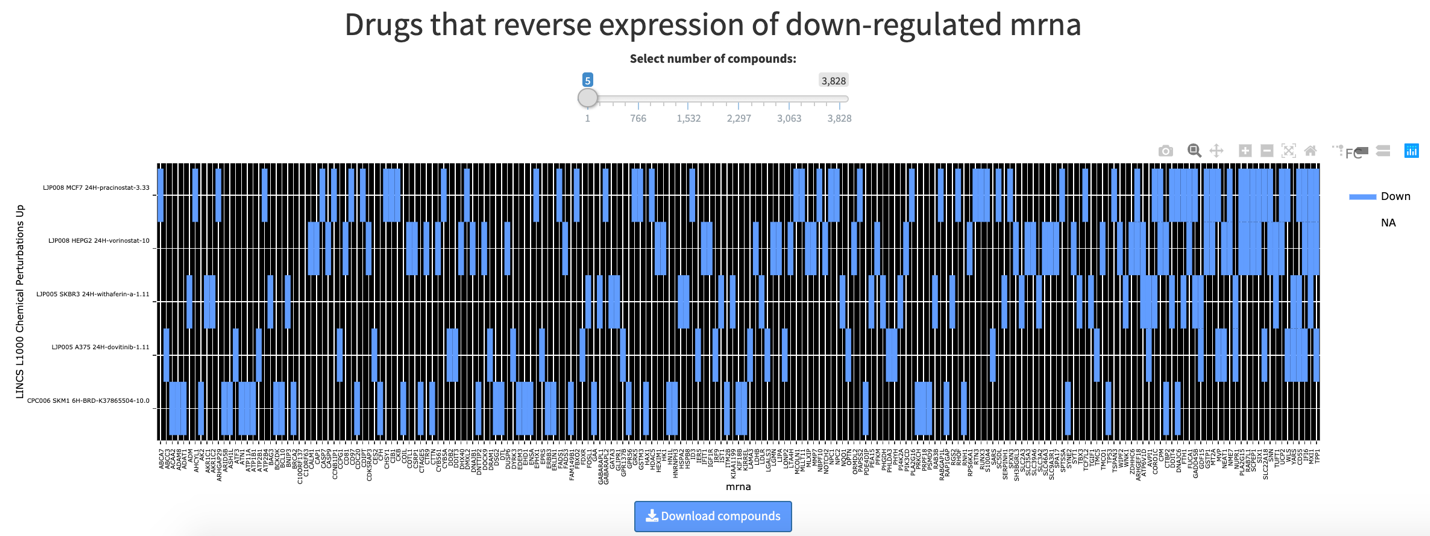

**B.**

**Supplementary Figure 4: Drug enrichment analysis using EnrichR.** For datasets in which the software detects gene symbols, gene set enrichment is automatically performed using the EnrichR API (Kuleshov *et al.*, 2016) using the LINCS_L1000_Chem_Pert_down and LINCS_L1000_Chem_Pert_down gene set collections based on the list of significant variables from Figure 2.The 1969 up- and 3145 down-regulated mRNA transcripts are used to identify chemical compounds that can reverse the expression profiles (from Figure 2). A. Chemical compounds that decrease the expression of a significant number of up-regulated genes by SARS-COV-2. B. Chemical compounds that increase the expression of a significant number of down-regulated genes by SARS-COV-2. Each plot can be zoomed in further to enable better viewing of compound and gene names.

**Case Study 2: Identify multi-omic biomarker panels of heart failure hospitalizations**

*Methods:* Blood samples was collected from 58 patients with heart failure and profiled for gene expression using microarrays (5000 gene transcripts) and protein expression using mass spectrometry (65 proteins). 29 electrical variables from Holter monitors were also collected. Cell-type frequencies were estimated using cell marker genes. These four omics datasets were used to identify a subset of biomarkers that were predictive of 3-month cardiac related hospitalizations. (See data compilation steps here: <https://github.com/singha53/omicsBioAnalytics/blob/master/inst/extdata/heartFailure/heartFailure.md>)

Steps:

1. **Data Upload**

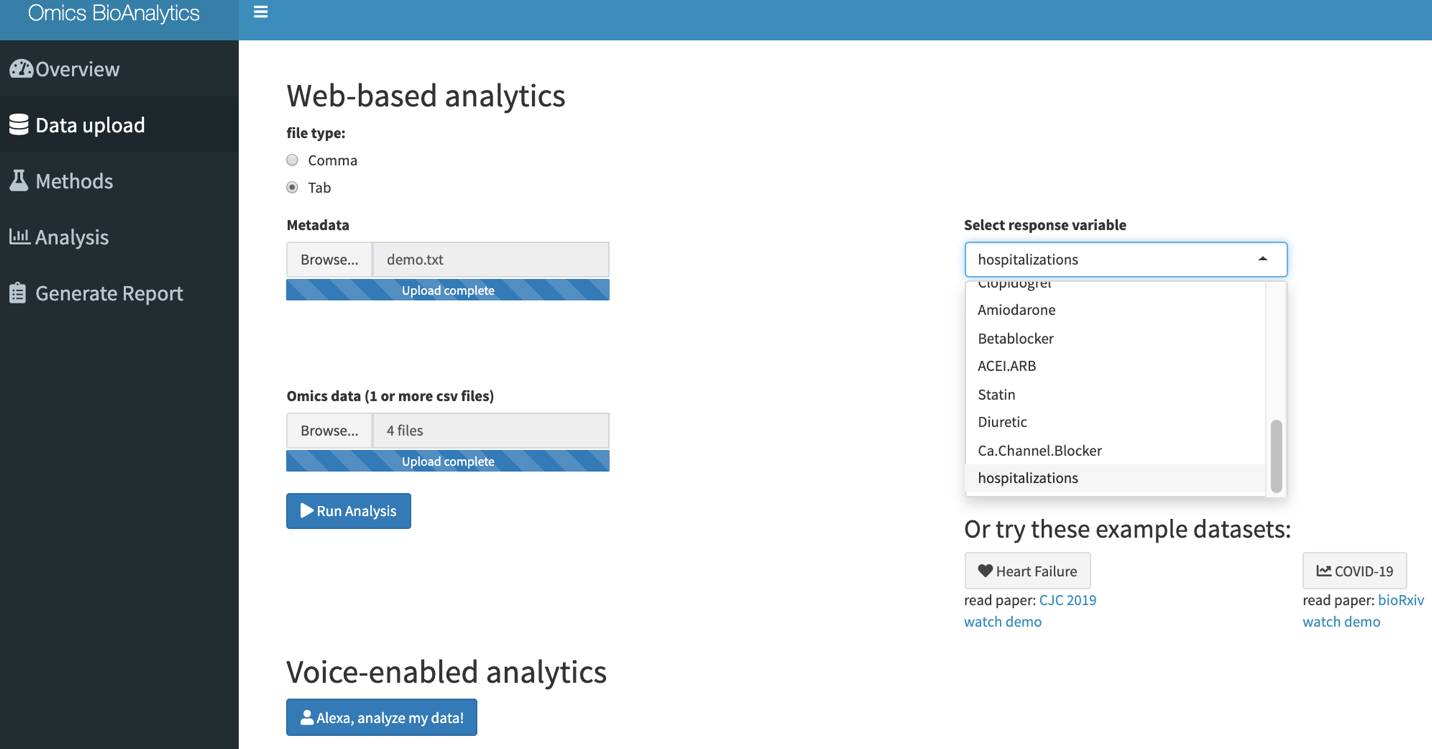

**Supplementary Figure 5. Data upload of heart failure datasets.** Metadata of the samples can be uploaded which prompts the user to select a response variable as well as a reference category. In this case, a demographics dataset was uploaded, and hospitalization status was chosen from the list of categorical variables with two or more categories. One or more omics datasets can also be uploaded, in this case four text files named *cells, holter*, *mrna,* and *proteins* were uploaded. All files have samples along the rows and variables along the columns.

2. **Metadata Analysis**

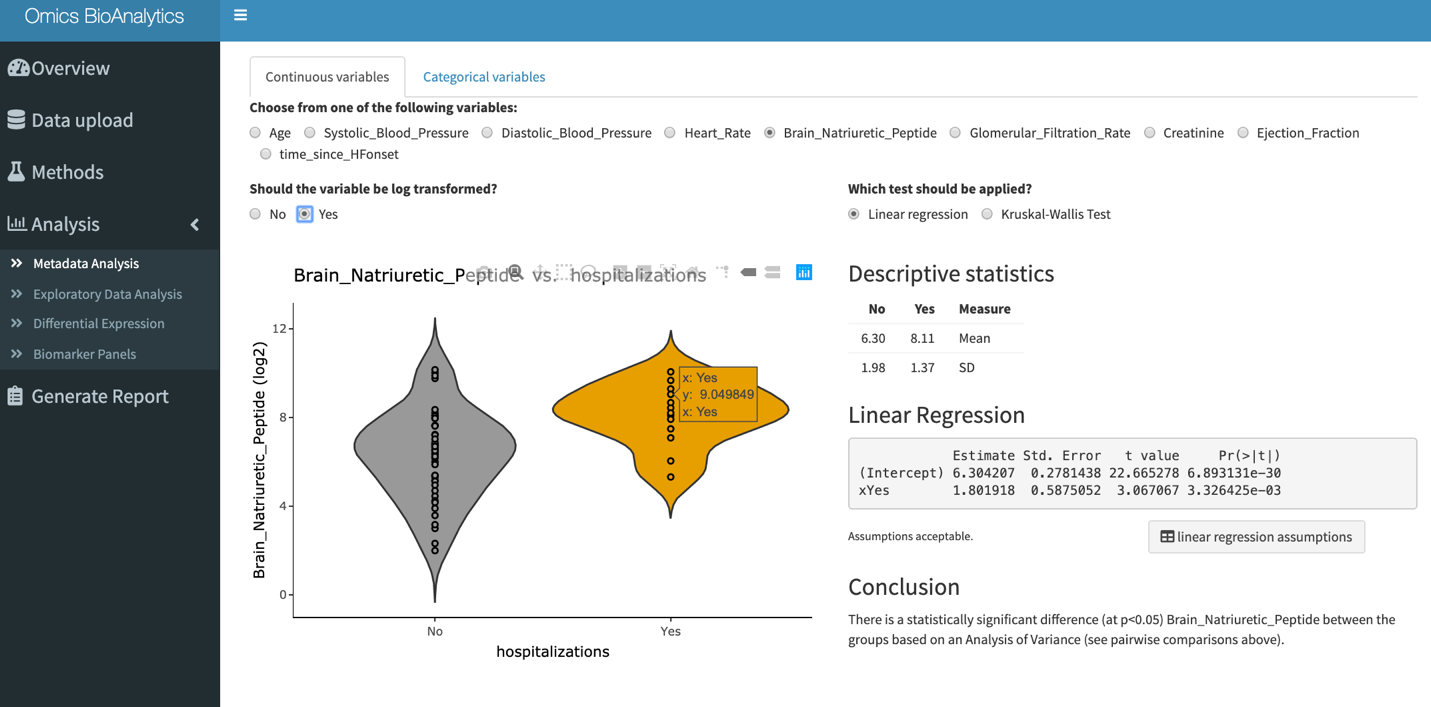

**Supplementary Figure 6. Analysis of continuous metadata variables.** Continuous variables are automatically identified from the metadata file and displayed as radio buttons for individual exploration. The example shows that the average levels of Brain Natriuretic Peptide (BNP) are significantly higher in patients that were hospitalized as compared to those that were not (p<0.05). The scale has been log2-transformed given the data is highly skewed.

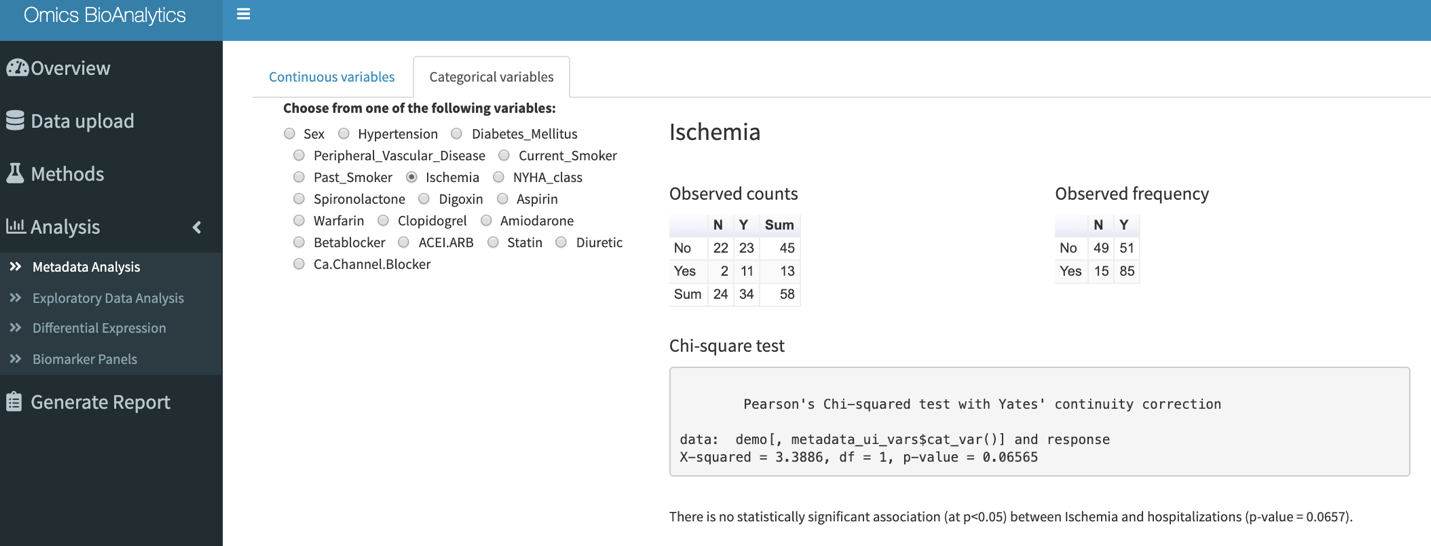

**Supplementary Figure 7. Analysis of categorical metadata variables.** Categorical variables are automatically identified from the metadata file and displayed as radio buttons for individual exploration. The example shows that contingency table hospitalization status and ischemia where no significant association between the observed and expected counts was identified (p=0.07)

3. **Exploratory Data Analysis**

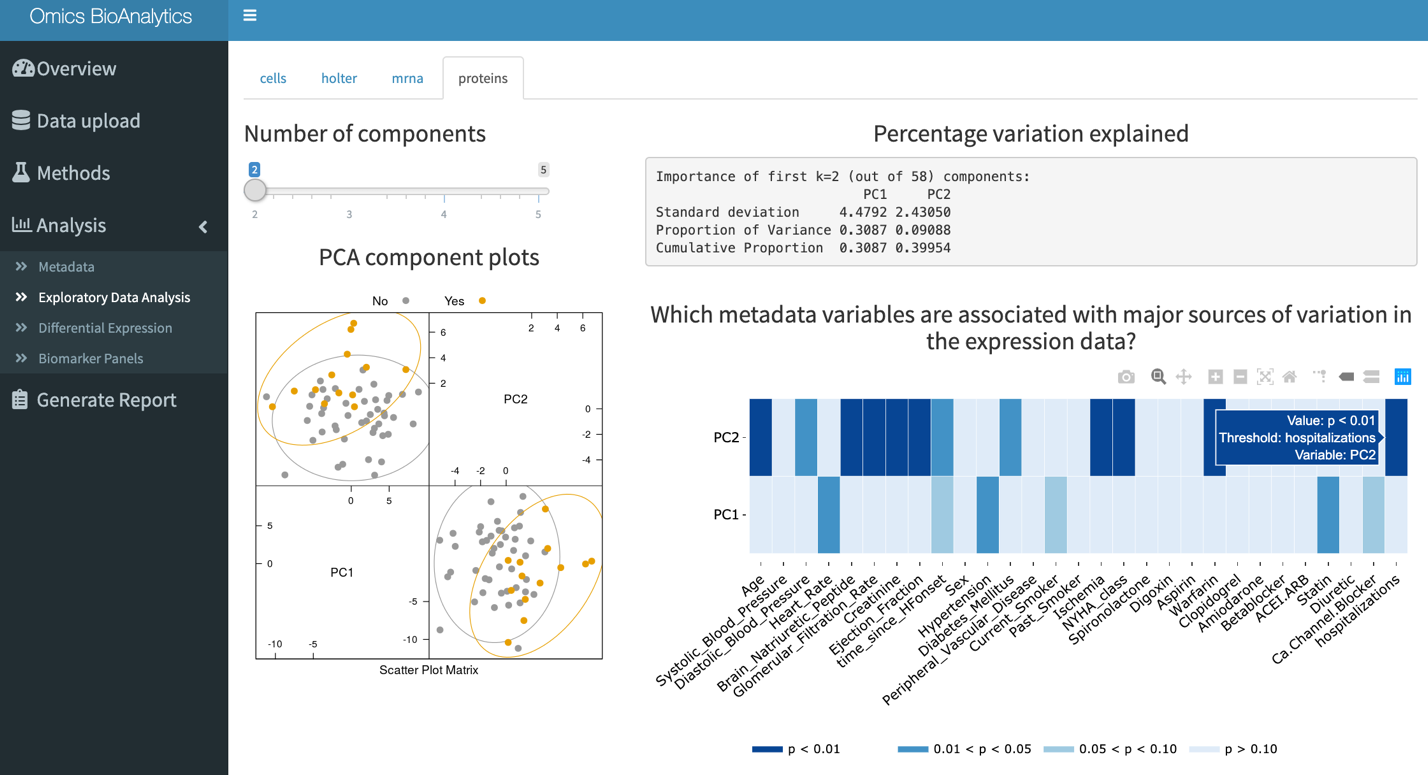

**Supplementary Figure 8. Exploratory data analysis of each uploaded omics dataset.** For each uploaded omics dataset principal component analysis (PCA) is performed. The user can select the number of principal components to depict in the scatter plot matrix. The heatmap depicts the ANOVA p-values (categorized into bins) comparing each principal component with each metadata variable. The figure above shows the PCA plot of the proteomics dataset for the first two principal components, colored by the response variable (hospitalization status) selected during the data upload stage. The heatmap shows the association of each metadata variable with a principal component. For example, hospitalization status is significantly associated with the second principal component (p<0.05), which can also be observed in the scatter plot matrix.

4. **Biomarker Discovery Analysis**

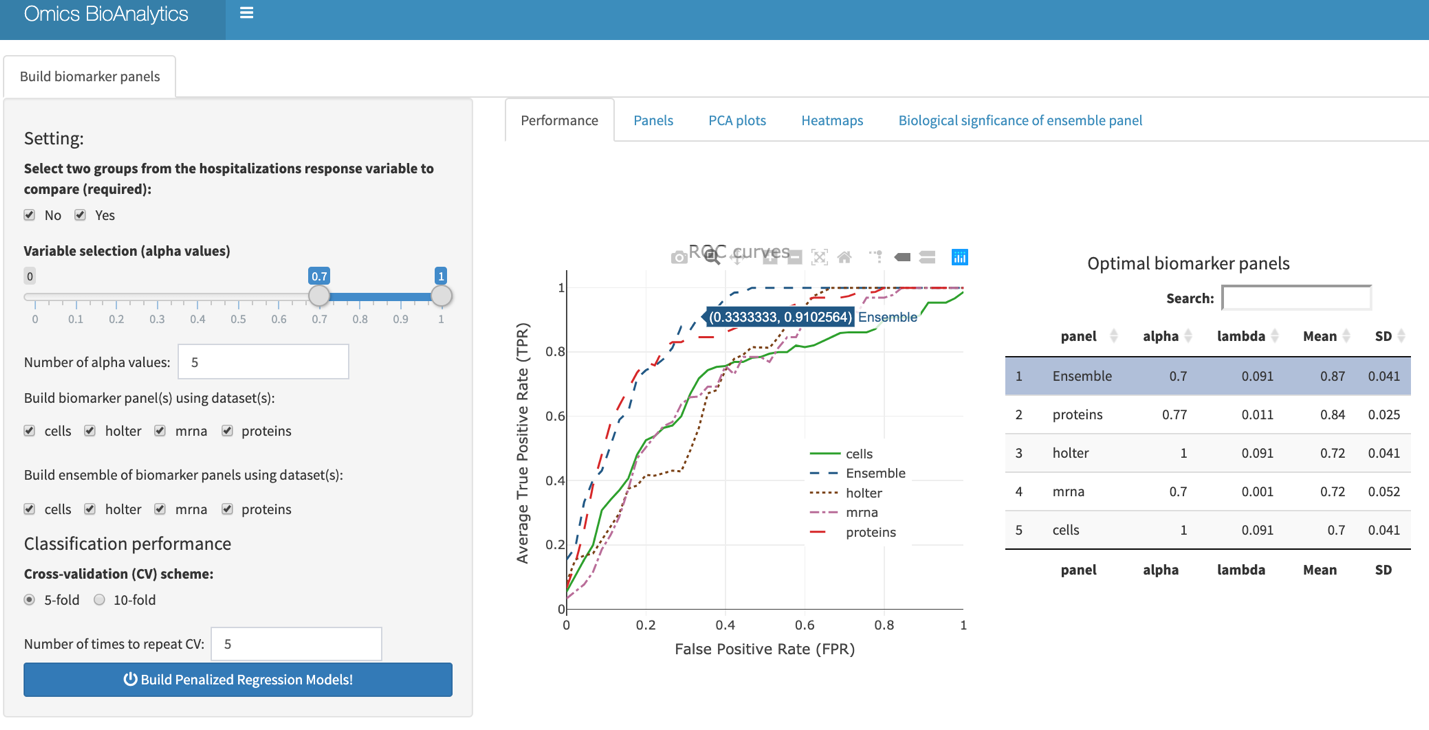

**Supplementary Figure 9. Biomarker discovery analysis – Performance tab.** Classification models using penalized regression models (glmnet R-library) are built for each omics dataset for a grid of hyperparameter values [for variable selection (alpha) and shrinkage of regression coefficients (lambda)]. The user can set a grid for the number of variables to select from each omics dataset [alpha=1: LASSO regression (few variables retained in final model), alpha=0: ridge regression (all variables retained in final model)]. The performance is evaluated using 5-fold or 10-fold cross-validation and repeated k times (set by user). The user can select which omics datasets to use to build classifiers for and which ones to combine in the ensemble model. The figure shows the selected datasets that were used to develop individual biomarker panels and the ensemble biomarker panel, the grid used for hyperparameter tuning includes five alpha values from 0.7 to 1 (0.700, 0.775, 0.850, 0.925, 1.000). The average receiver operating characteristic (ROC) curves are depicted with the corresponding summary table, showing that the Ensemble biomarker panel outperforms the individual biomarker panels with an AUC=0.87.

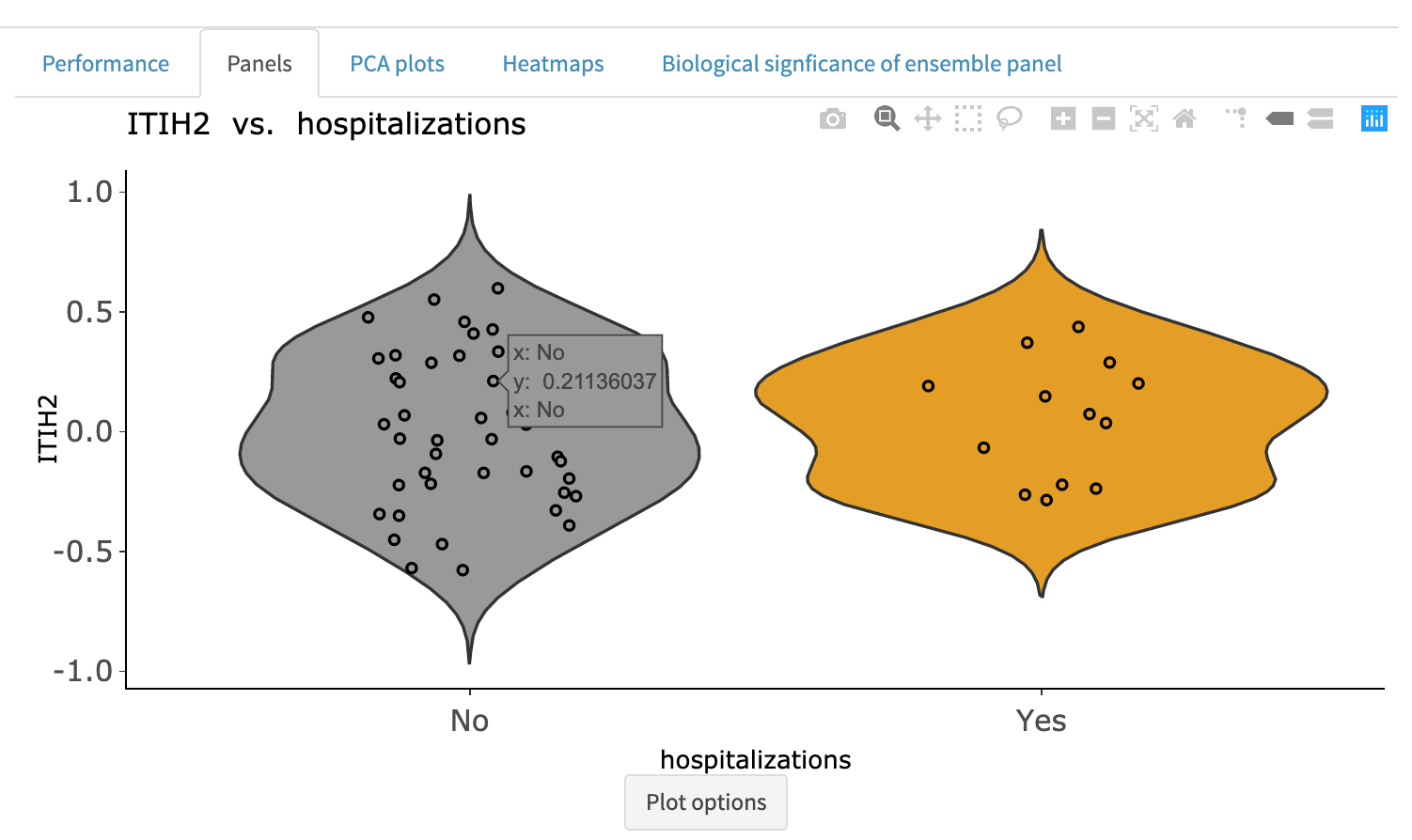

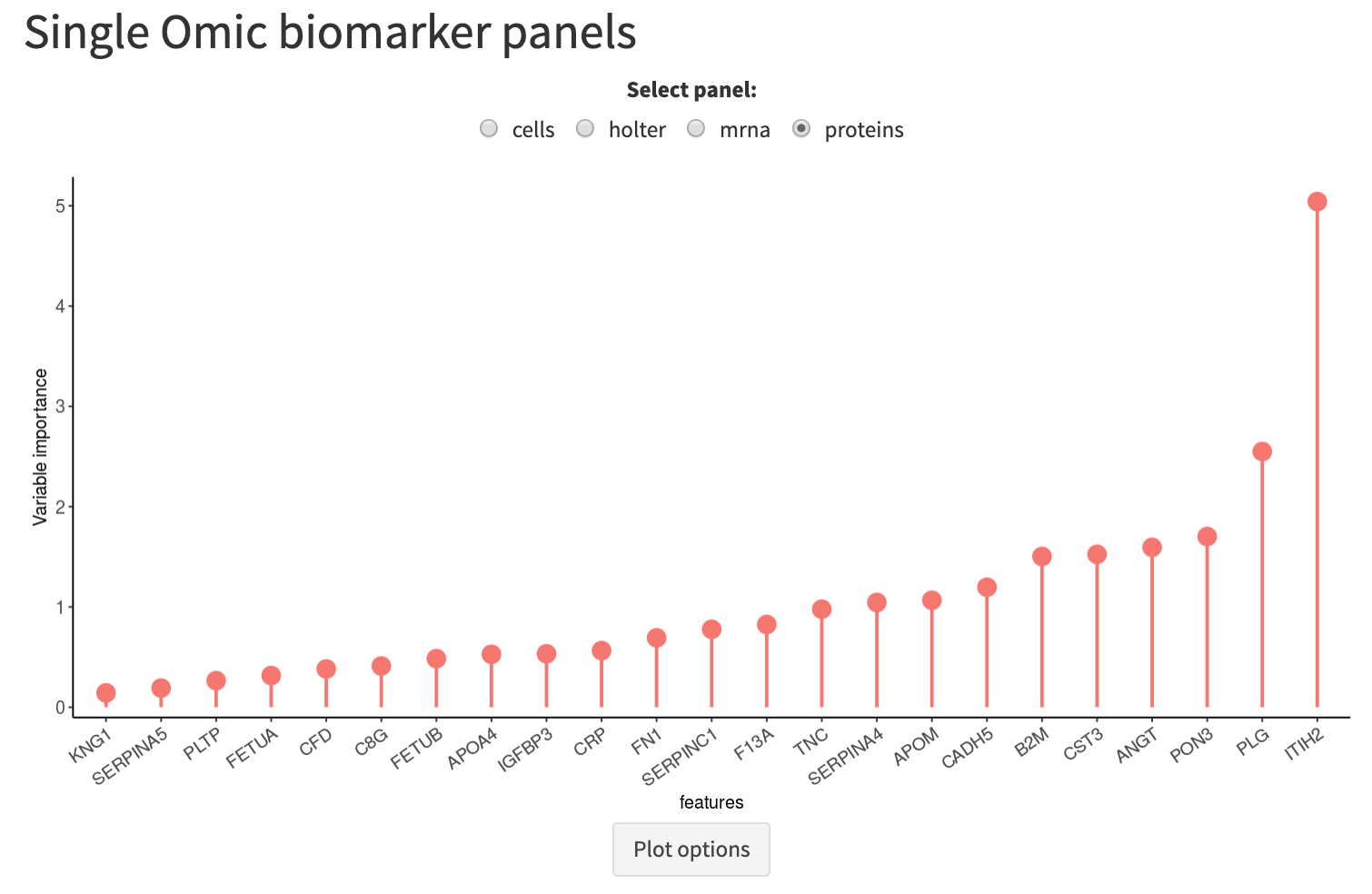

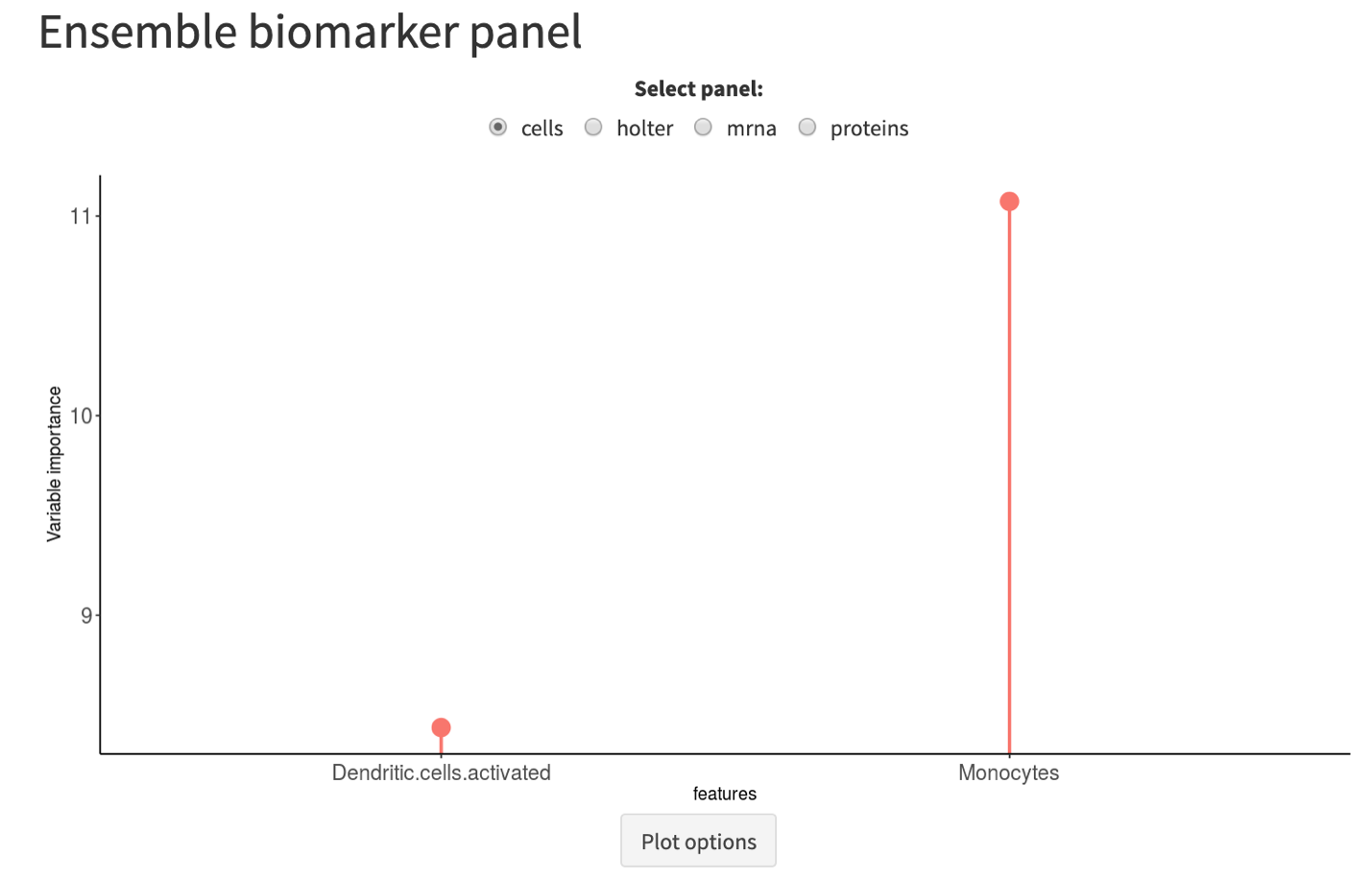

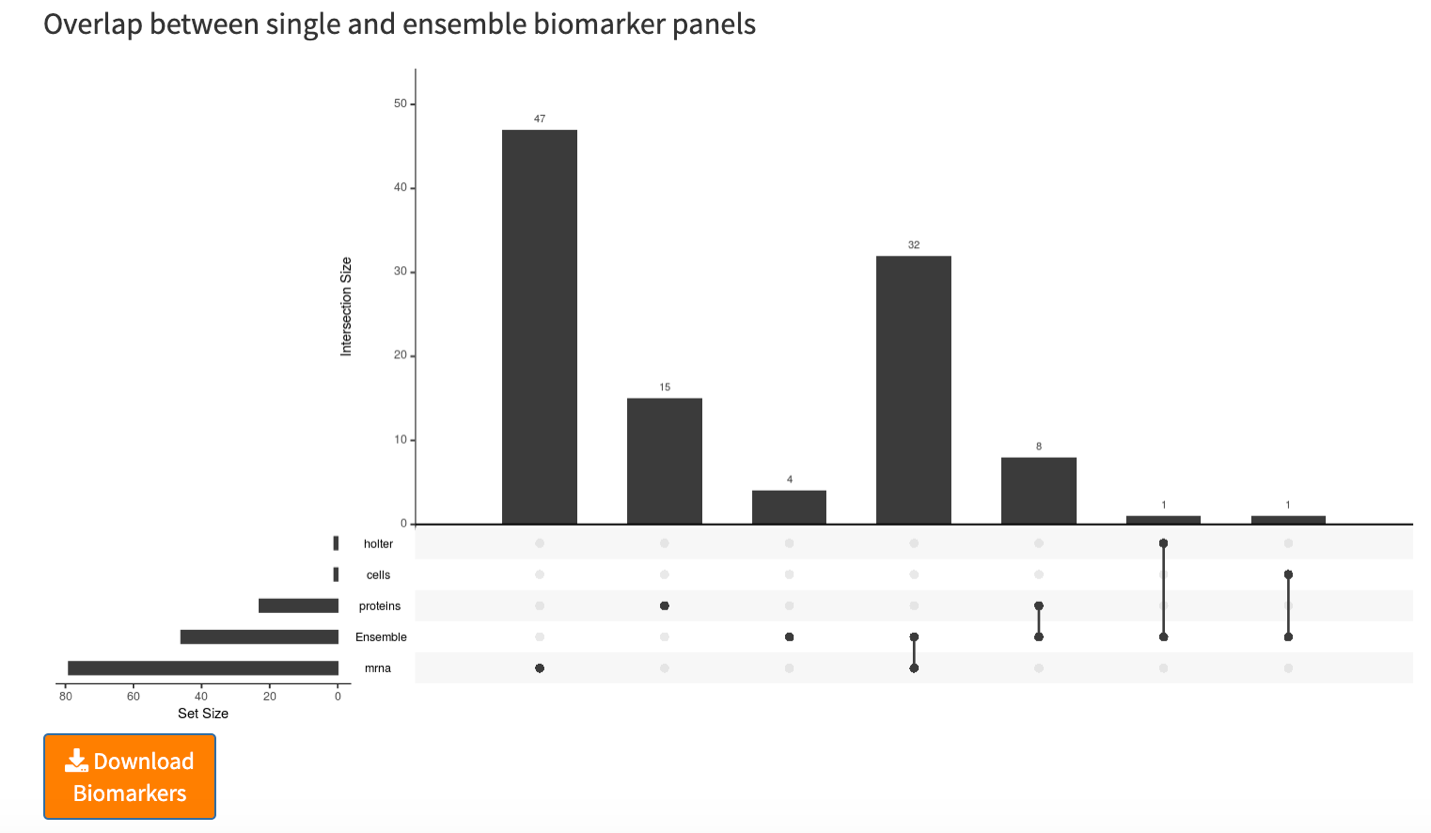

**Supplementary Figure 10.** **Biomarker discovery analysis – Panels tab.** Selected variables in the individual biomarker panels and ensemble biomarker panel are depicted using dot plots. Clicking on a given dot will display the specific variable, in this example, ITIH2 of the protein biomarker panel was clicked and the associated violin plot is displayed. The intersection plot displays the number of overlaps between each pair of panels, and in this example, the greatest overlap is between the ensemble biomarker panel and the mRNA biomarker panel. The list of all biomarkers can also be downloaded.

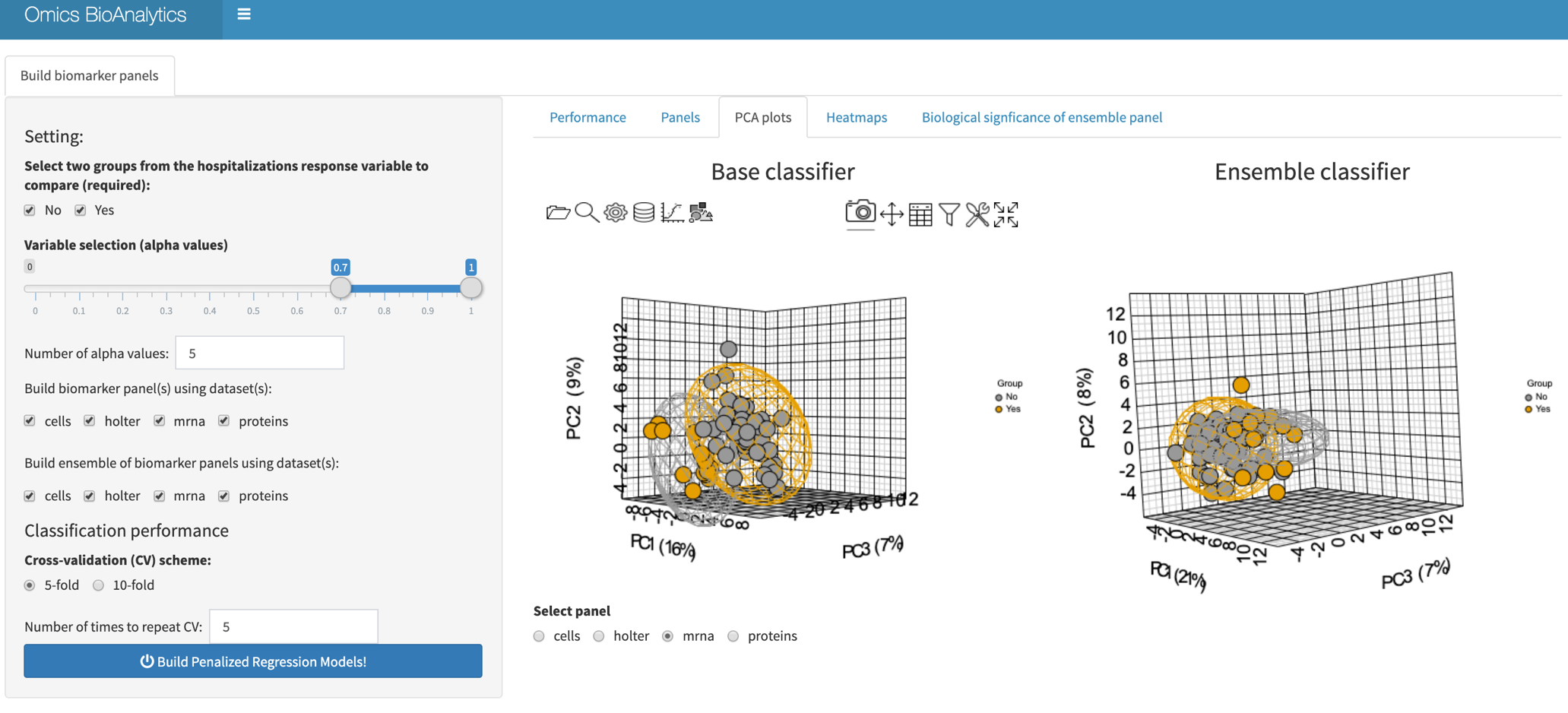

**Supplementary Figure 11.** **Biomarker discovery analysis – PCA plots tab.** The figure displays the PCA plots using biomarkers in the mRNA base classifier (left) and using biomarkers of various data types (cells, holter, mRNA and protein) in the ensemble classifier (right).

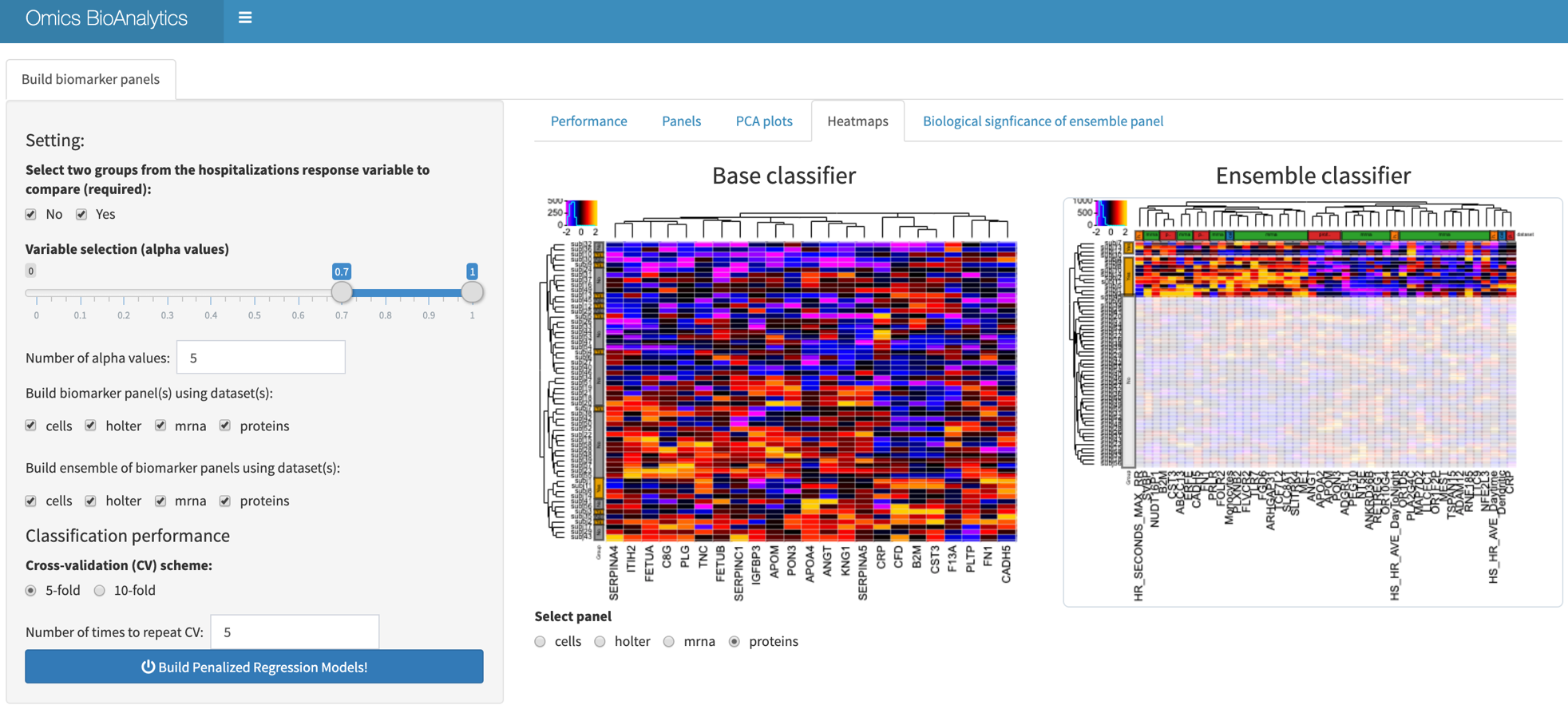

**Supplementary Figure 12.** **Biomarker discovery analysis – Heatmaps tab.** The hierarchical clustering of samples and variables based on the protein biomarkers is displayed using a heatmap on the left whereas biomarkers of various data types (cells, holter, mrna and protein) in the ensemble classifier is displayed on the right. Data is centered and scaled and capped at ±2 prior to plotting. Clicking on the row or column labels of the heatmaps can be used to narrow in on the clustering patterns of specific subgroups or data types. For example, for the ensemble classifier, rows corresponding to samples labelled with a hospitalization status of *Yes* are highlighted.

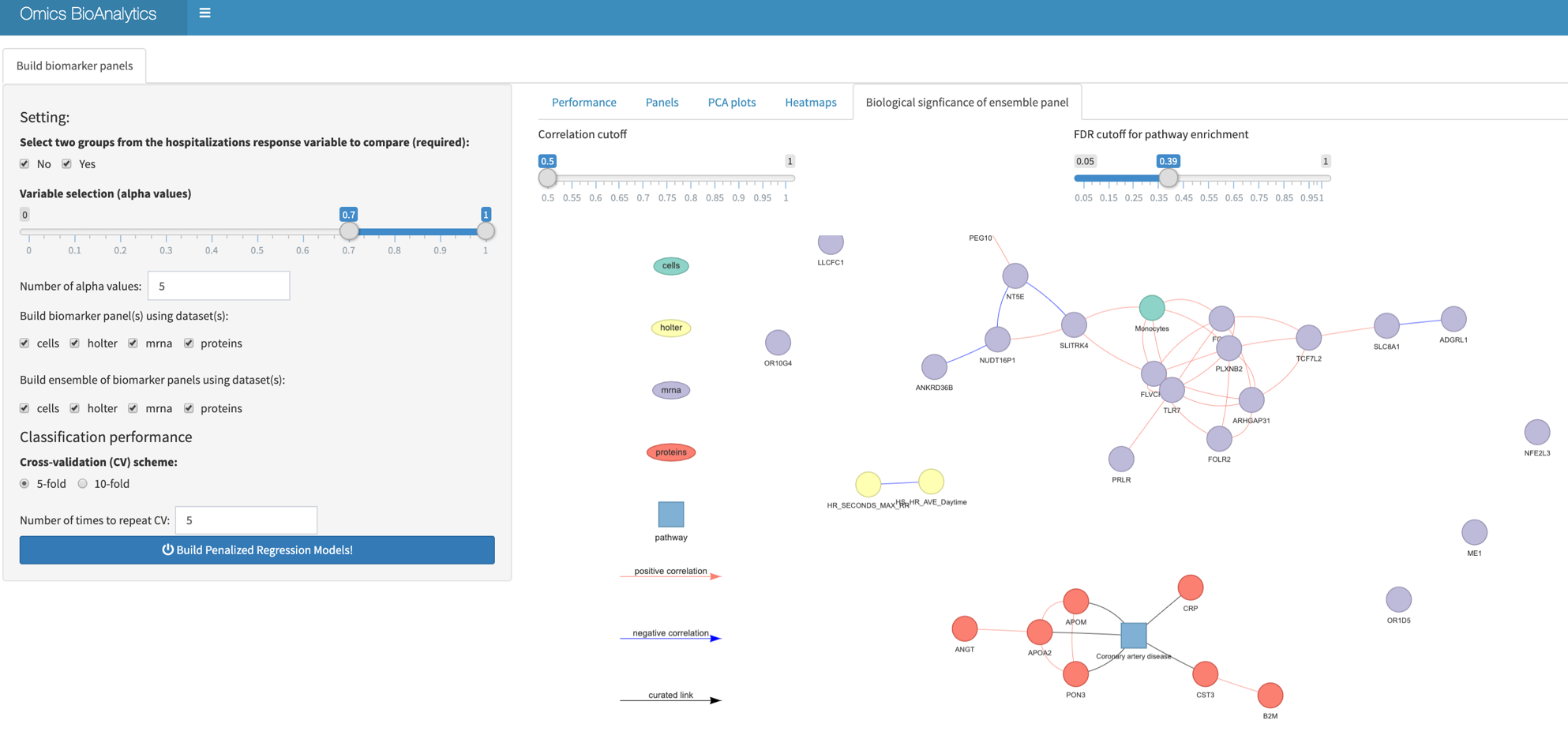

**Supplementary Figure 13.** **Biomarker discovery analysis – Biological significance of ensemble panel tab.** Enrichment analysis of variables in the ensemble biomarker panel is performed using various thresholds for significance (geneset enrichment analysis) and pairwise correlations. The sub-network on the bottom connected proteins (APOM, APOA2, PON3, CRP and CST3) to curated pathway *Coronary artery disease* and this network was further extended by including other correlated proteins (ANGT and B2M).
